# Redundant and additive functions of the four Lef/Tcf transcription factors in lung epithelial progenitors

**DOI:** 10.1101/2020.01.29.925842

**Authors:** Kamryn N. Gerner-Mauro, Haruhiko Akiyama, Jichao Chen

## Abstract

In multicellular organisms, paralogs from gene duplication survive purifying selection by evolving tissue-specific expression and function. Whether this genetic redundancy is also selected for within a single cell type is unclear for multi-member paralogs, as exemplified by the 4 obligatory Lef/Tcf transcription factors of canonical Wnt signaling, mainly due to the dauntingly complex genetics involved. Using the developing mouse lung as a model system, we generate 2 quadruple conditional knockouts and myriad triple and double combinations, and show that the 4 Lef/Tcf genes function redundantly in the presence of at least 2 Lef/Tcf paralogs, but additively upon losing additional paralogs to specify and maintain lung epithelial progenitors. Pre-lung-specification, pan-epithelial double knockouts have no lung phenotype, triple knockouts have varying phenotypes, including defective branching and tracheoesophageal fistulas, and the quadruple knockout barely forms a lung, resembling the *Ctnnb1* mutant. Post-lung-specification deletion of all 4 Lef/Tcf genes leads to branching defects, downregulation of progenitor genes, premature alveolar differentiation, and derepression of gastrointestinal genes, again phenocopying the corresponding *Ctnnb1* mutant. Our study supports a monotonic, positive signaling relationship between CTNNB1 and Lef/Tcf in lung epithelial progenitors and represents, to our knowledge, the first in vivo analysis of cell-type-specific genetic redundancy among the 4 Lef/Tcf paralogs.

**SIGNIFICANCE STATEMENT:** Paralogs represent genetic redundancy and survive purifying selection by evolving overlapping and distinct functions. In multicellular organisms, such functional diversification can manifest as tissue and cell type specific expression, which masks possible selective pressure for genetic redundancy within a single cell type. Using in vivo genetic and genomic analyses, we show that although the 4 mammalian Lef/Tcf transcription factors have evolved organ-specific functions, they function additively and redundantly, depending on gene dosage, to promote lung epithelial progenitors and do so in a monotonic, positive manner with beta-Catenin in the canonical Wnt signaling pathway.

## INTRODUCTION

Sequenced genomes of key species in the phylogenetic tree, especially the amphioxus, suggest that 2 rounds of whole genome duplication quadruple ancestral genes in the jawed vertebrates, as exemplified by the single Hox gene cluster in flies and 4 in mammals (1–3). This, together with individual gene amplification, generates paralogs that are initially redundant and must evolve unique functions to escape purifying selection. This apparent genetic redundancy is maintained if the surviving paralogs provide (i) an extra amount of the same function, (ii) a back-up to buffer environmental fluctuations including somatic mutations, or (iii) a new function such as alternative enzymatic substrates (4). In multicellular organisms, when paralogs differ in their tissue and cell type distributions, such differential expression even in the absence of differential protein activity provides unique functions and allows their existence in the genome. This organism-level genetic redundancy, however, masks possible redundancy within individual cell types, which is only revealed by studying cell-type-specific mutants.

The canonical Wnt signaling pathway culminates in beta-Catenin (CTNNB1) binding to and activating the Lef/Tcf transcription factors, which include a single member in flies (Pangolin or dTCF) and worms (POP-1) but 4 in mice and humans (LEF1, TCF7, TCF7L1, and TCF7L2) (5). Global, single knockout of individual Lef/Tcf genes in mice results in defects in skin appendages (*Lef1*), T cells (*Tcf7*), embryonic axis induction (*Tcf7l1*), and intestinal epithelial stem cells (*Tcf7l2*), highlighting the aforementioned tissue-specific functions of paralogs. Several subsequent double knockout models demonstrate their overlapping but non-interchangeable functions, and partially phenocopy deletion of their obligatory co-factor CTNNB1 (6). Studying additional paralogs involves dauntingly complex genetic crosses and is limited to cultured cells and global deletion in early Xenopus embryos (7, 8). As a result, genetic redundancy of Lef/Tcf genes in vivo and within a given cell type, as well as the corresponding requirement of CTNNB1, is unknown.

The established requirement of WNT2/2B and CTNNB1 in specifying lung epithelial progenitors from the embryonic foregut, as well as the continued role of CTNNB1 in maintaining the progenitors (9–11), supports the involvement of canonical Wnt signaling and lays the foundation to investigate the identity and associated genetic redundancy of Lef/Tcf transcription factors. In this study, using pan-lung-epithelium and progenitor-specific Cre drivers, we first deleted *Tcf7l1* and *Tcf7l2*, the main Lef/Tcf paralogs expressed in the lung epithelium, but did not detect any phenotype. Extending our analysis to additional double mutants, all possible triple mutants, and the quadruple mutant, we show that depending on gene dosage, Lef/Tcf paralogs function additively and redundantly in lung progenitor specification and branching. Single cell RNA-seq (scRNA-seq) reveals the role of Lef/Tcf paralogs in maintaining lung progenitors by promoting progenitor genes and suppressing gastrointestinal genes and premature alveolar differentiation, recapitulating the transcriptional defects of the progenitor-specific *Ctnnb1* mutant. Our study clarifies the signaling relationship between CTNNB1 and Lef/Tcf and provides an in vivo demonstration of genetic redundancy within a single cell type.

## RESULTS

### TCF7L1 and TCF7L2, the main LEF/TCF factors in the lung epithelium, are not required for epithelial progenitor specification and maintenance

The lung epithelial primordium originates from the anterior foregut as a group of NKX2-1/SOX9 double positive progenitors, which remain distally during subsequent branching morphogenesis, leaving behind SOX2-expressing progeny for the airways early in development and alveolar cells late in development (12). The established role of canonical Wnt signaling components, CTNNB1 and WNT2/2B, in specifying and maintaining these lung progenitors (9–11) provides an in vivo system to investigate the 4 corresponding transcription factor paralogs: LEF1, TCF7, TCF7L1, and TCF7L2 (6). We set out to identify which LEF/TCF was present in the epithelium using immunostaining, and found that while all 4 factors were expressed in the surrounding mesenchyme, only TCF7L1 and TCF7L2 were readily detectible in the epithelium (Fig. 1A). Accordingly, we deleted both *Tcf7l1* and *Tcf7l2* using *Shh*^*Cre*^ (13) throughout the epithelium and before lung specification (Fig. 1B). In contrast to the lung agenesis phenotype of the pan-epithelial *Ctnnb1* mutant (9–11), the *Tcf7l1*/*Tcf7l2* double mutant lungs had normal branching structure and branch morphology with normal distal and proximal distributions of SOX9 and SOX2, respectively (Fig. 1C). Furthermore, alveolar differentiation, as measured by HOPX for alveolar type 1 (AT1) cells and LAMP3 for alveolar type 2 (AT2) cells, was also unaffected (Fig. 1C).

**Figure 1.**
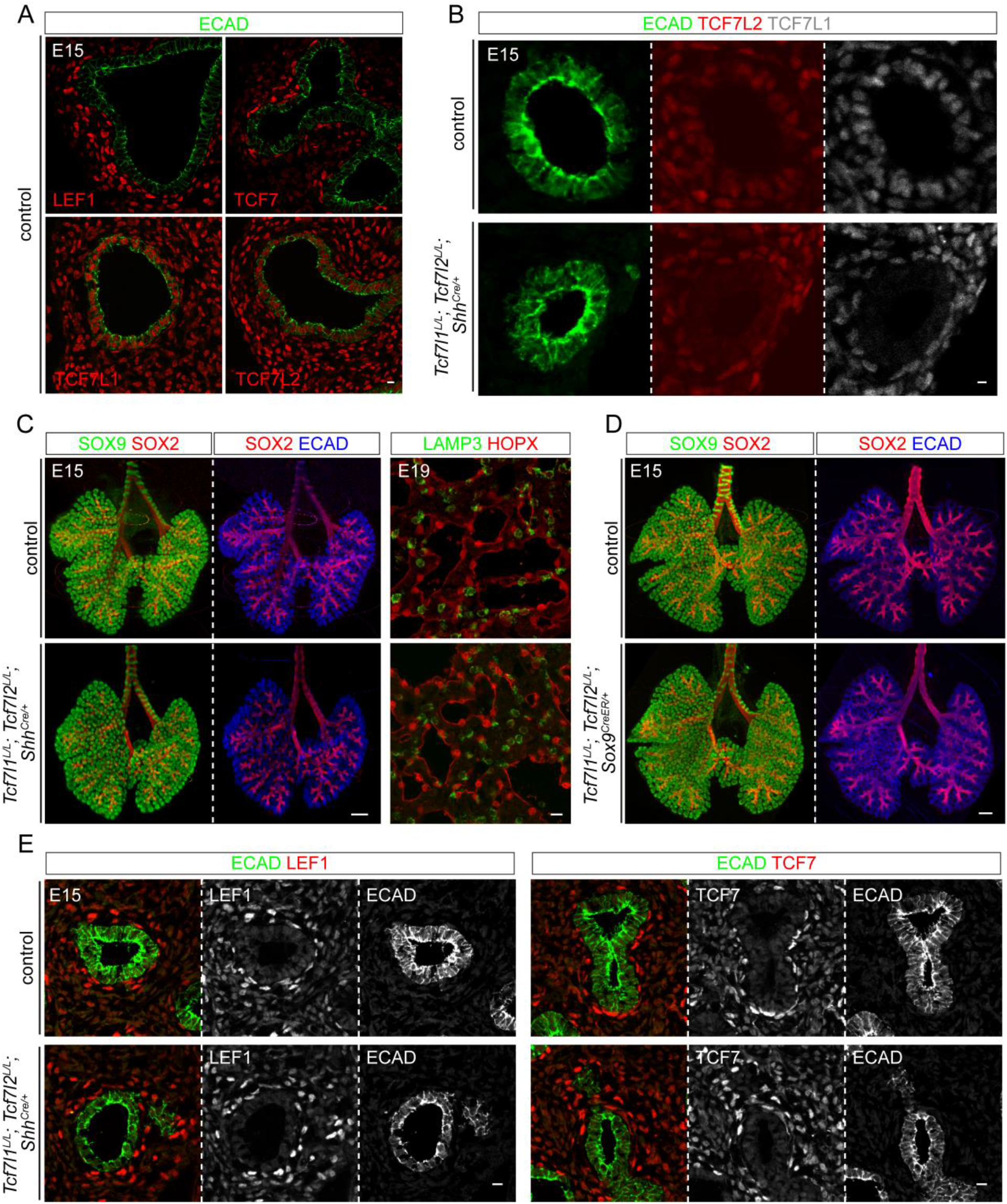
TCF7L1 and TCF7L2, the main LEF/TCF factors in the lung epithelium, are not required for epithelial progenitor specification and maintenance. **(A)** Confocal images of immunostained control lungs showing expression of all 4 Lef/Tcf factors in the mesenchyme, but mostly TCF7L1 and TCF7L2 in the epithelium outlined by ECAD. **(B)** Confocal images of immunostained littermate control and *Tcf7l1*^*L/L*^; *Tcf7l2*^*L/L*^; *Shh*^*Cre/+*^ mutant lungs showing loss of TCF7L1 and TCF7L2 in the mutant epithelium. **(C)** Optical projection tomography (OPT) images (left 2 column) and confocal images (rightmost column) of immunostained littermate control and *Tcf7l1*^*L/L*^; *Tcf7l2*^*L/L*^; *Shh*^*Cre/+*^ mutant lungs showing no difference in branching morphogenesis and distribution of SOX9 and SOX2, nor difference in AT1 (HOPX) and AT2 (LAMP3) cell differentiation. **(D)** OPT images of immunostained littermate control and *Tcf7l1*^*L/L*^; *Tcf7l2*^*L/L*^; *Sox9*^*CreER/+*^ mutant lungs showing no difference in branching. **(E)** Confocal images of immunostained littermate lungs showing no compensatory upregulation of LEF1 or TCF7. All images are representative of at least 2 biological replicates. Scale bars: 10 µm for confocal images and 250 µm for OPT images.

To account for any transient phenotype that might be compensated and thus masked in the pan-epithelial, pre-lung-specification *Shh*^*Cre*^ mutant (14), we also deleted *Tcf7l1* and *Tcf7l2* using a progenitor-specific driver *Sox9*^*CreER*^ and induced recombination at embryonic day (E) 11, right after lung specification; we still did not observe any morphological or molecular change (Fig. 1D). We formally examined possible compensatory upregulation of LEF1 and TCF7 in the pan-epithelial *Tcf7l1*/*Tcf7l2* double mutant, and found that they remained barely detectible (Fig. 1E). These data showed that despite their dominant expression in the epithelium, TCF7L1 and TCF7L2 are not required for lung progenitor specification and maintenance.

### LEF1/TCF7/TCF7L1/TCF7L2 function additively and redundantly to specify epithelial progenitors

The lack of phenotype in the *Tcf7l1*/*Tcf7l2* double mutant raised the possibility that LEF1 and TCF7 may be functional without being expressed at an appreciable level. Consistent with this possibility, all 4 LEF/TCF proteins were detected in the epithelium at E10 when the lung buds just emerged from the foregut (Fig. 2A). Since expression level might not be predictive of function (Fig. 1), we resorted to genetics to generate various combinations of double, triple, and quadruple Lef/Tcf mutants using *Shh*^*Cre*^. Like the *Tcf7l1*/*Tcf7l2* double mutant, none of the other double mutants had a lung phenotype, as assayed by morphology and distribution of distal (SOX9) and proximal (SOX2) markers (Fig. 2B). We did not generate the *Lef1/Tcf7l2* double mutant because, as detailed below, even deleting one more paralog in 2 of the triple mutants did not result in any phenotype.

**Figure 2.**
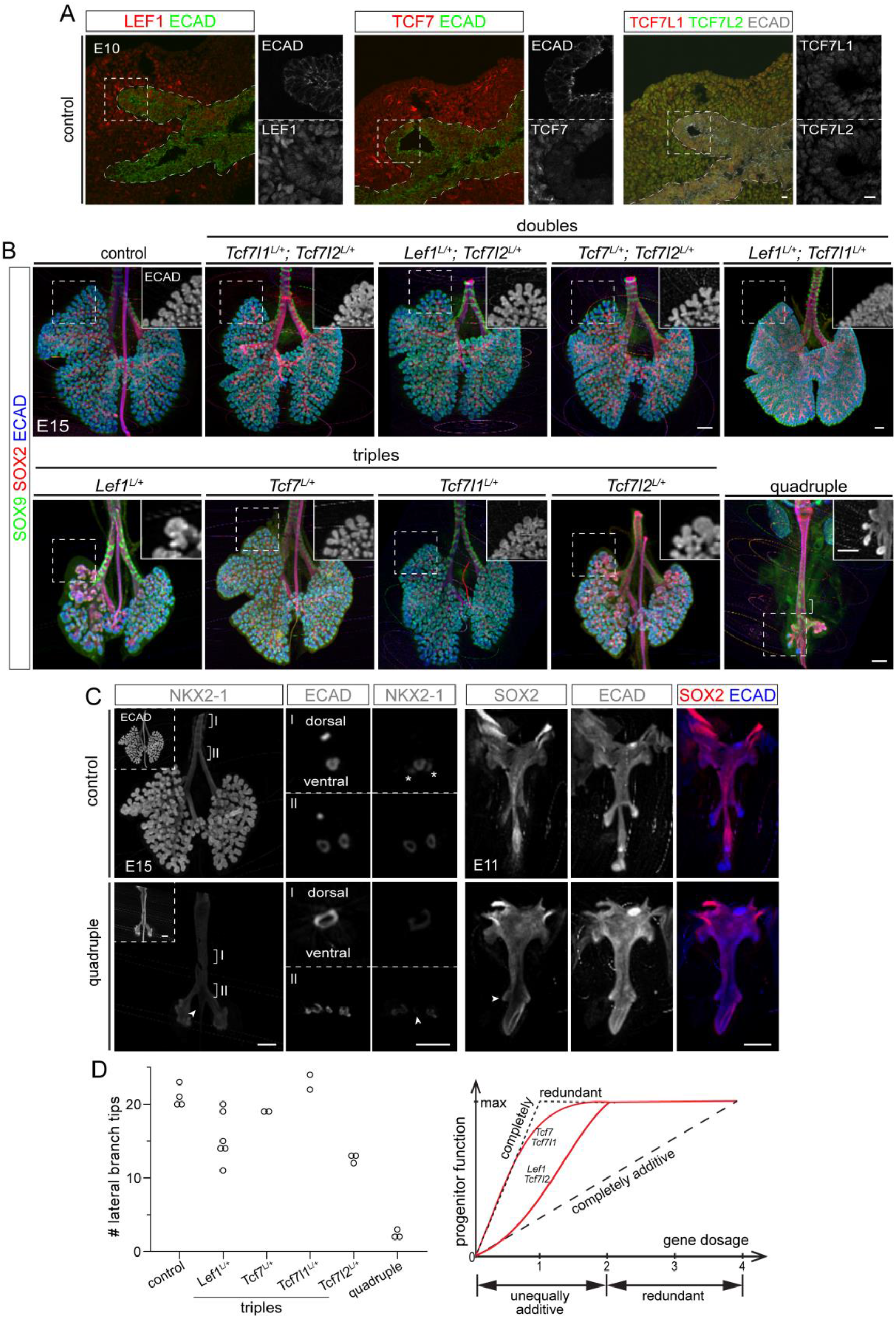
LEF1/TCF7/TCF7L1/TCF7L2 function additively and redundantly to specify lung epithelial progenitors. **(A)** Confocal images of immunostained control lungs at the time of specification showing all 4 Lef/Tcf factors detectible in the epithelium. **(B)** OPT images of immunostained control, double, triple, and quadruple mutants. All mutants are epithelial specific using *Shh*^*Cre*^. Six out of 8 alleles in the double mutants are conditional with the remaining 2 wild type alleles as indicated; they all have normal branching. Seven out of 8 alleles in the triple mutants are conditional with the remaining wild type allele as indicated; they have varying phenotypes as detailed in the main text. Boxed regions are shown as insets for ECAD staining. The square bracket in the quadruple mutant indicates SOX9 positive cartilage rings on the ventral side of the fused foregut tube. See also supplemental videos 1-6. **(C)** OPT images of immunostained littermate control and quadruple mutant lungs. Left: the mutant has reduced NKX2-1, which is enriched on the ventral side and extends aberrantly beyond the bifurcation point of left and right main bronchi (arrowhead). Numbered brackets indicate regions shown as optical sections. Asterisks, non-specific NKX2-1 staining outside the epithelium. Right: the mutant has aberrant branches as early as E11 and also aberrant distal expression of SOX2 (arrowhead). See also supplemental videos 7-8. **(D)** Quantification of lateral branch tips of the left lobes of control, triple, and quadruple mutant lungs at E15. Each symbol represent one lung. The 2 seemingly normal *Lef1*^*L/+*^ triple mutants are defective in branching (Fig. S1D). **(E)** Schematics illustrating that Lef/Tcf act redundantly with 2 or more paralogs present and additively albeit unequally upon further deletion (red lines), in comparison with theoretical modes of completely redundant and completely additive (black dash). All images are representative of at least 2 biological replicates. Scale bars: 10 µm for confocal images and 250 µm for OPT images.

Among the 4 triple mutants where 7 out of 8 alleles were targeted by *Shh*^*Cre*^, the *Lef1*/*Tcf7l1*/*Tcf7l2* and *Lef1*/*Tcf7*/*Tcf7l2* mutants had normal branching (Fig. 2B), whereas the remaining 2 triple mutants had smaller lungs with fewer and dilated branch tips, characteristic of defective branching morphogenesis of the lung (12) (Fig. 2B, Videos 1-5). Although still expressing SOX9, these abnormal branches often had SOX2 expression extended further distally and, at times, completely replacing SOX9 expression, suggesting thwarted SOX9 progenitor expansion and/or their accelerated differentiation into SOX2 progeny (Fig. 2B). The *Tcf7*/*Tcf7l1*/*Tcf7l2* triple mutant additionally had a fused trachea and esophagus – recapitulating the tracheoesophageal fistula phenotype in the pan-epithelial *Ctnnb1* mutant (9–11) (Fig. 2B, Fig. S1A). Intriguingly, whereas no viable pups were obtained for the 3 triple mutants carrying *Tcf7l2* deletion presumably due to the reported intestinal defects (15), the *Lef1*/*Tcf7*/*Tcf7l1* triple mutant – despite defective branching – formed apparently normal alveoli and survived postnatally, albeit with no hair as expected from *Lef1* deletion (16) (Fig. S1B). These data suggest that when only 1 of the 4 Lef/Tcf paralogs remains in the lung progenitors, individual paralogs are no longer redundant nor equivalent: TCF7 or TCF7L1 supports sufficient Wnt signaling whereas LEF1 or TCF7L2 is partially functional.

Finally, the *Shh*^*Cre*^ quadruple mutant lung consisted of 2 small sacs that fused with the esophagus and expressed SOX2 except for occasional clusters of SOX9 cells that might have escaped recombination and represented failed attempts to branch due to their limited number (Fig. 2B, Fig. S1C, Video 6). Furthermore, the expression level of the lung lineage transcription factor NKX2-1 was noticeably reduced albeit still enriched in the ventral portion of the fused tracheoesophageal tube, suggesting defective lung fate specification (Fig. 2C). In addition, NKX2-1 expression tapered off beyond the aberrant lung sacs along the ventral side of the esophagus tube, possibly due to an anterior shift in lung specification or ectopic posterior specification by WNT2/2B-expressing mesenchyme that failed to extend away as a result of defective branching. The reduction in NKX2-1 was also observed in our published *Ctnnb1* mutant where *Ctnnb1* is deleted shortly after lung specification at E11 (11), but different from the complete absence of NKX2-1 in the pre-lung-specification *Ctnnb1* mutant or the *Wnt2/2b* mutant (9, 10). A possible explanation is that deletion of all 8 alleles is not fast enough by a single Cre driver and occurs shortly after lung specification. The branching phenotype could be traced back to E11 when the left and right lung buds in the quadruple mutant failed to extend and were covered by SOX2 expression (Fig. 2C, Videos 7, 8).

The graded phenotype of the triple and quadruple mutants suggested an additive function among the Lef/Tcf paralogs and was quantified by counting lateral branch tips in the left lobe (17) (Fig. 3D, Fig. S1D, Videos 1-6). This allowed us to propose a model of Lef/Tcf function in the lung progenitors (Fig. 3E): when 2 or more paralogs are present, they act redundantly and at a full level of Wnt signaling; when only 1 paralog is present, Wnt signaling can fall below a threshold with individual paralogs functioning additively but unequally; and there is minimal Wnt signaling when all 4 paralogs are missing. This provided direct, in vivo evidence for the selective advantage of maintaining genetic redundancy of 4 paralogs within a single cell type. The triple mutants also revealed the function of individual Lef/Tcf paralogs and supported a monotonic, positive relationship between Lef/Tcf and CTNNB1 in the lung progenitors, different from the proposed repressor function in other contexts (18–21).

**Figure 3.**
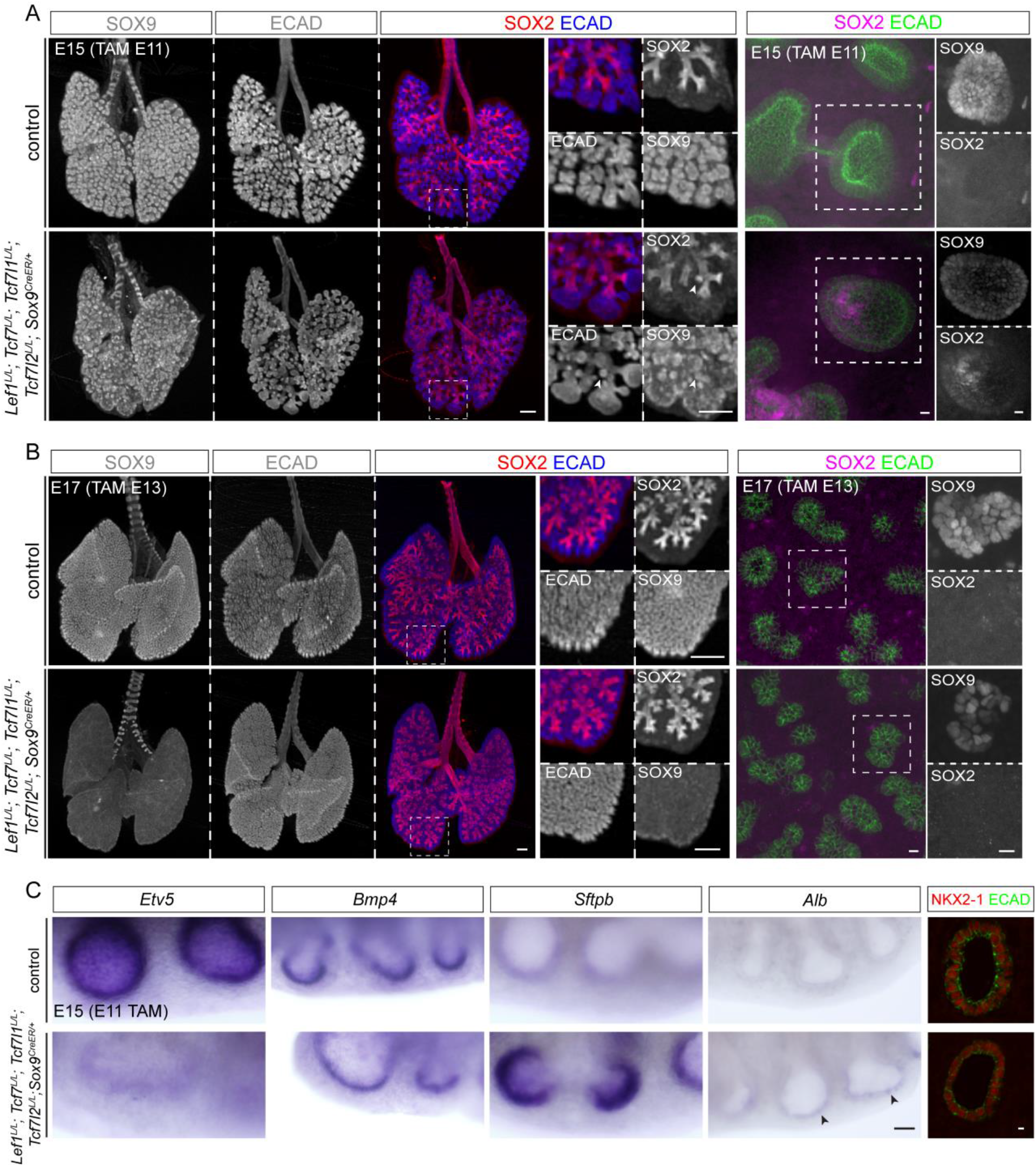
LEF1/TCF7/TCF7L1/TCF7L2 maintain lung epithelial progenitors. (**A**, **B**) OPT images (left) and confocal image stacks of cranial lobes (right) of littermate control and post-lung-specification (*Sox9*^*CreER*^) quadruple mutant lungs at indicated time points. E11 deletion (**A**) leads to branch dilation, SOX9 reduction, and distal extension of SOX2 (arrowhead). E13 deletion (**B**) leads to SOX9 reduction, but no apparent change in branch morphology or SOX2 expression. TAM: tamoxifen. See also supplemental videos 9-10. (**C**) Whole mount in situ hybridization (left) and confocal section immunostaining (right) of littermate control and *Sox9*^*CreER*^ quadruple mutant lungs showing reduction of a Wnt-dependent progenitor gene *Etv5*, premature expression of an alveolar gene *Sftpb*, and ectopic expression of a gastrointestinal gene *Alb* (arrowheads), but no change in a Wnt-independent progenitor gene *Bmp4* or NKX2-1. All images are representative of at least 2 biological replicates. Scale bars: 10 µm for confocal images and 250 µm for OPT and in situ images.

### LEF1/TCF7/TCF7L1/TCF7L2 maintain epithelial progenitors

Since the *Shh*^*Cre*^ quadruple mutant barely formed a lung, we sought to characterize the function of Lef/Tcf in the progenitors by deleting all 4 paralogs with *Sox9*^*CreER*^ after lung specification, as we have done to study CTNNB1 (11). When recombination was induced at E11, progenitor branching was compromised, as reflected in the characteristic dilation of branch tips, in association with downregulation of SOX9 (Fig. 3A, Videos 9, 10). Some branch tips had SOX2 expression over-extended distally, suggesting an imbalance favoring airway differentiation over progenitor expansion (Fig. 3A). When recombination was induced at E13, SOX9 was similarly downregulated, although branch morphology and SOX2 expression were unaffected at E17 (Fig. 3B), possibly because functional loss of Lef/Tcf paralogs occurred after the progenitors had initiated alveolar differentiation.

These morphological and molecular changes largely phenocopied the post-lung-specification *Ctnnb1* mutant (11). Such similarity was reinforced by the same downregulation of additional, but not all, progenitor genes (as exemplified by *Etv5* and *Bmp4*, respectively), premature expression of an alveolar differentiation gene *Sftpb*, and ectopic expression of a gastrointestinal gene *Alb* (Fig. 3C)(11). However, unlike the *Ctnnb1* mutant, NKX2-1 expression was unaffected even when Lef/Tcf recombination was induced early at E11 (Fig. 3C). This might be because it took longer to recombine all 8 alleles when NKX2-1 expression no longer depended on Wnt signaling, as supported by its normal expression when *Ctnnb1* was deleted at E13 (11). Taken together, Lef/Tcf paralogs, similar to CTNNB1, maintain lung epithelial progenitors by promoting a subset of progenitor genes and repressing genes associated with alveolar and gastrointestinal differentiation.

### Single-cell RNA-seq profiling of post-lung-specification Lef/Tcf mutants reveal ectopic gastrointestinal differentiation and premature alveolar differentiation

To delineate the transcriptional program that depended on Lef/Tcf paralogs, we preformed scRNA-seq of E15 *Sox9*^*CreER*^ quadruple mutants with recombination induced at E11 and their littermate controls. To identify changes in the epithelium, we used our published fluorescence-activated cell sorting protocol to separate the immune, endothelial, epithelial, and mesenchymal cells using CD45, ICAM2, and ECAD (Fig. 4A) (22). A total of 2,071 and 4,062 epithelial cells from control and quadruple mutant lungs, respectively, were sequenced; both genotypes consisted of 3 epithelial cell types – SOX9 progenitors, SOX2 progeny, and neuroendocrine cells, the last of which were marked by *Ascl1* and were among the first to mature in the lung epithelium (23) (Fig. 4B). A lower proportion of SOX9 progenitors was present in the Lef/Tcf mutant lung (68% versus 43%), consistent with the observed expansion of SOX2 immunostaining (Fig. 3A). In the reduced dimension plot, only the SOX9 progenitor population was shifted between control and mutant lungs, indicating significant transcriptional changes specifically in SOX9 progenitors (Fig. 4B). This was supported by differential expression analysis of each of the 3 cell types: nearly all differentially expressed genes, including a Wnt target gene *Axin2* (24), were in SOX9 progenitors, except for a few for the SOX2 progeny population that were likely due to perdurance of SOX9 progenitor genes during airway differentiation (Fig. 4C, Table S1). Corroborating and extending our in situ hybridization analysis of candidate genes (Fig. 3C), downregulated genes included some but not all progenitor genes, such as *Sox9*, *Etv5*, *Lin7a*, and *Wnt7b*; upregulated genes included markers of alveolar differentiation, such as *Igfbp7* and *Sftpb*, and those of gastrointestinal differentiation, such as *Alb*, *Gkn2*, and *Apoa1* (Fig. 4C, Table S1).

**Figure 4.**
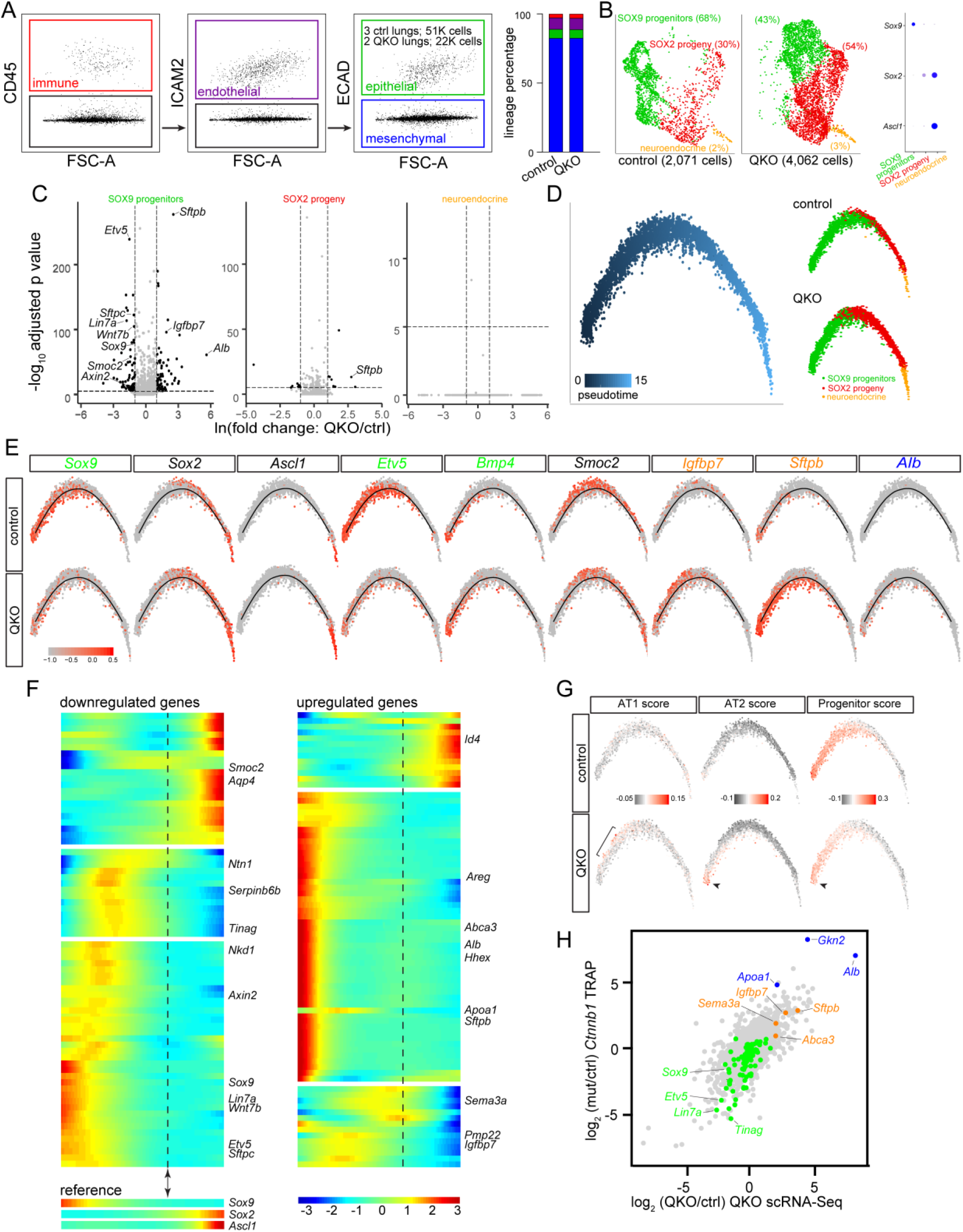
Single-cell RNA-seq profiling of the post-lung-specification (*Sox9*^*CreER*^) Lef/Tcf mutant reveals ectopic gastrointestinal differentiation and premature alveolar differentiation. **(A)** Left: FACS gating strategy to purify lung epithelial cells. 3 littermate control lungs yielded 50,000 epithelial cells and 2 quadruple mutant lungs yielded 22,000 epithelial cells. Right: percentages of lung lineages from FACS, color-coded as in the left panels. QKO (quadruple knockout): *Lef1*^*L/L*^; *Tcf7*^*L/L*^; *Tcf7l1*^*L/L*^; *Tcf7l2*^*L/L*^; *Sox9*^*CreER/+*^ at E15 with tamoxifen administered at E11. **(B)** Seurat Uniform Manifold Approximation and Projection for Dimension Reduction (UMAP) plots after excluding non-epithelial cells showing 3 epithelial cell types with the corresponding percentages in parenthesis, as identified by marker genes in the dot plot. **(C)** Volcano plots comparing control and mutant lungs for the SOX9, SOX2, and neuroendocrine populations. See also Table S1. **(D)** Monocle trajectory of control and mutant lungs, color-coded by cell type, transitions from SOX9 progenitor to SOX2 progeny and neuroendocrine cells. **(E)** Distributions of cell population markers and differentially expressed genes along the Monocle trajectory. Green: progenitor genes including Wnt-dependent *Sox9* and *Etv5* as well as Wnt-independent *Bmp4*. Orange: alveolar genes including *Igfbp7* (AT1) and *Sftpb* (AT2). Blue: gastrointestinal genes including *Alb*. *Smoc2* is expressed in cells transitioning from SOX9 to SOX2 and is lower in the mutant. **(F)** Monocle branch heatmaps of downregulated and upregulated genes from the SOX9 volcano plot. The SOX9-SOX2 transition is marked by a double arrowhead and vertical dashes, based on reference gene expression. See Table S2 for complete gene lists. **(G)** Distributions of metagene scores using signature genes of AT1, AT2, and progenitor cells. AT1 gene signature is upregulated past the trajectory origin (bracket), whereas upregulation of AT2 gene signature and downregulation of progenitor gene signature occur at the trajectory origin (arrowheads). The remaining progenitor score in the mutant is driven by Wnt-independent progenitor genes. See Table S3 for gene lists. Top 100 AT1 and AT2 genes are obtained from an E19 scRNA-seq dataset (Table S4). **(H)** Comparison of quadruple mutant scRNA-seq and *Ctnnb1* mutant bulk RNA-seq, color-coded as in (**E**). See also Table S5.

To capture the transcriptional dynamics accompanying active cell differentiation and maturation, we performed Monocle trajectory analysis and found that SOX9 progenitors, SOX2 progeny, and neuroendocrine cells formed a linear trajectory, consistent with *Sox9*^*CreER*^ lineage-tracing experiments (17, 25) (Fig. 4D). As expected, progenitor genes were highest at the origin of the trajectory and a subset of them were lower in the Tcf/Lef mutant (Fig. 4E, F). Notably, upregulated genes including those of alveolar and gastrointestinal differentiation were also biased toward the trajectory origin, suggesting that such aberrant activation occurred only in SOX9 progenitors but was somehow suppressed as mutant cells differentiated into their SOX2 progeny (Fig. 4E, F). Interestingly, some genes including *Sema3a*, *Pmp22*, and *Igfbp7* were activated in mutant cells right after the trajectory origin and were normally enriched in alveolar type 1 (AT1) cells, whereas genes of alveolar type 2 (AT2) cells including *Sftpb* and *Abca3* were activated at the origin (Fig. 4E, F and Table S2). These changes were confirmed in a metagene analysis using AT1 and AT2 gene signatures derived from scRNA-seq of E19 lungs (26), as well as our published 119 progenitor genes (11) (Fig. 4G, Table S3, Table S4). Such trajectory bias of AT1 and AT2 genes in this premature alveolar differentiation was reminiscent of their spatial bias during normal alveolar differentiation (27, 28). Finally, transcriptomic changes in the quadruple mutant mirrored those induced by *Ctnnb1* deletion (11) (Fig. 4H, Table S5). Taken together, our transcriptomic analysis, together with our genetic evidence, supports a positive functional correlation between Lef/Tcf paralogs and CTNNB1.

### DISCUSSION

In this study, we have generated 2 conditional quadruple mutants and used 3D imaging and single-cell genomics to define the in vivo function of the 4 Lef/Tcf paralogs of the canonical Wnt signaling pathway. We demonstrate that the 4 Lef/Tcf paralogs function additively and redundantly to specific lung epithelial progenitors: 2 paralogs in any combination or TCF7 or TCF7L1 alone are fully functional, whereas LEF1 or TCF7L2 alone is partially functional and losing all paralogs phenocopies losing CTNNB1, the canonical Wnt signaling cofactor. Our study reinforces the concept of lung progenitors balancing dynamic competing cell fates, provides in vivo evidence for a monotonic positive relationship between Lef/Tcf paralogs and CTNNB1, and sheds light on genetic redundancy.

The lung epithelial progenitors expand from a few hundred when specified from the embryonic foregut to millions within a week of branching morphogenesis. Such dramatic progenitor expansion is even more remarkable considering that they have the potential to undergo airway, alveolar, or even gastrointestinal differentiation (11, 12). While gastrointestinal differentiation is likely undesirable in the lung, the plasticity in choosing airway and alveolar fates and the timing of doing so might provide an opportunity to evolve lungs of different complexity, as suggested by the primitive, 2-sac frog lungs (29). By pinpointing the identity of the 4 Lef/Tcf paralogs involved, this study supports 2 pivotal roles of canonical Wnt signaling in regulating the lung progenitors: (i) promoting progenitor branching while delaying alveolar differentiation, and (ii) preventing gastrointestinal differentiation. Future work using low-cell-number and single-cell approaches will reveal the associated direct transcriptional targets and the normal temporal shift in progenitor potentials, thereby establishing an epigenetic roadmap of lung progenitor development and the corresponding transcriptional regulatory network. Such knowledge is essential for deriving human lung cells in culture for both mechanistic and therapeutic research, especially given the pro-airway and pro-gastrointestine effects of inhibiting Wnt signaling (30).

In the most intuitive model, canonical Wnt signaling culminates in CTNNB1 binding and activation of Lef/Tcf transcription factors, such that CTNNB1 and Lef/Tcf function in a monotonic, positive manner. Nevertheless, extensive research – as warranted by deployment of Wnt signaling in numerous biological contexts – suggests that some Lef/Tcf paralogs, in particular TCF7L1, could also function as a transcriptional repressor (31). However, since all Lef/Tcf paralogs can biochemically interact with transcriptional corepressors (20), overexpression experiments might introduce non-physiological activities. Adding to the complexity, available Wnt reporters differ significantly, including in the lung (32). In the current study, the established role of CTNNB1 allows us to ascertain the function of individual Lef/Tcf paralogs by comparing the corresponding triple mutants with the quadruple mutant (Fig. 2). Specifically, TCF7 or TCF7L1 alone is sufficient to transmit CTNNB1 function, while LEF1 or TCF7L2 is partially effective, suggesting all Lef/Tcf paralogs are transcriptional activators in lung epithelial progenitors, although future work should examine whether CTNNB1 and Lef/Tcf paralogs directly or indirectly suppress alveolar and gastrointestinal differentiation. Intriguingly, as in lung epithelial progenitors, TCF7 or TCF7L1 is also sufficient to support embryonic stem cell differentiation (8). Extending these comprehensive genetic deletion analyses to other cell types is necessary to clarify the transcriptional activity of the Lef/Tcf paralogs.

Our study provides in vivo evidence for genetic redundancy in a specific cell type. Since all 4 Lef/Tcf paralogs are present in the fish, predating the appearance of the lung, it is conceivable that when Wnt signaling was coopted to evolve lung epithelial progenitors, a core transcriptional module common to all Lef/Tcf paralogs was used to allow their co-expression, generating the initial redundancy. Such redundancy presumably buffers somatic mutations and thereby confers a selective advantage given the obvious requirement of the lung for survival on land. Intriguingly, our triple mutants suggest that partial Wnt signaling supported by LEF1 or TCF7L2 alone leads to an underdeveloped lung that, although compatible with life at least in the case of TCF7L2, likely reduces the fitness and is selected against, reinforcing the importance of maintaining genetic redundancy. Nevertheless, epithelial expression of LEF1 and TCF7 is lower than their mesenchymal expression (Fig. 1A), suggesting possible relaxation of the selection pressure to maintain all 4 paralogs – perhaps an evolutionary intermediate toward more limited redundancy among fewer paralogs, as in more ancient systems such as the thymocytes for LEF1 and TCF7 (33). Such drift on the transcriptional level must, however, not interfere with expression elsewhere in the body; a systematic comparison of the underlying enhancers across cell types and species should shed light on the evolution of genetic redundancy.

## METHODS

### Mice (*Mus musculus*)

The following mouse strains were used: *Lef1*^*LoxP*^ (34), *Tcf7*^*LoxP*^ (35), *Tcf7l1*^*LoxP*^ (36), *Tcf7l2*^*LoxP*^ (15), *Shh*^*Cre*^ (13), and *Sox9*^*CreER*^ (37). The day of the presence of a vaginal plug was marked as embryonic day (E) 1. To induce Cre recombination, 3 mg of tamoxifen (T5648, Sigma) dissolved in corn oil (C8267, Sigma) was injected intraperitoneally at E11 or E13. All animal experiment were approved by the institutional animal care and use committee at MD Anderson Cancer Center.

### Antibodies

The following antibodies were used: rabbit anti-aquaporin 5 (AQP5, 1:250, ab78486, Abcam), PE/Cy7-conjugated rat anti-CD45 (CD45, 1:250, 103114, BioLegend), rat anti-E-cadherin (ECAD, 1:2000, 131900, Invitrogen), Alexa488-conjugated rat anti-E-cadherin (ECAD, 1:250, 53-3249-80, eBioscience), rabbit anti-Hopx (HOPX, 1:500, Santa Cruz, sc-30216), A647-conjugated rat anti-intercellular adhesion molecule 2 (ICAM2, 1:250, A15452, Invitrogen), eFluor570-conjugated rat anti-Mki67 (KI67, 1:500, 41-5698-82, eBioscience), rat anti-lysosomal-associated membrane protein 3 (LAMP3, 1:1000, DDX0192, Imgenex), rabbit anti-lymphoid enhancer binding factor 1 (LEF1, 1:500, 2230, Cell Signaling), rabbit anti-NK2 homeobox 1 (NKX2-1, 1:1000, sc-13040, Santa Cruz Biotechnology), goat anti-SRY-box-containing gene 9 (SOX9, 1:1000, AF3075, R&D systems), rabbit anti-SRY-box-containing gene 9 (SOX9, 1:1000, AB5535, Millipore), goat anti-SRY-box-containing gene 2 (SOX2, 1:250, sc-17320, Santa Cruz Biotechnology), rabbit anti-T cell factor 7 (TCF7, 1:250, 2203, Cell Signaling), guinea pig anti-T cell factor 7 like 1 (TCF7L1, 1:500, a gift from Dr. Hoang Nguyen at Baylor College of Medicine), and rabbit anti-T cell factor 7 like 2 (TCF7L2, 1:100, 2569, Cell Signaling).

### Section and whole-mount immunostaining

Immunostaining was performed as previously published with minor modifications (11, 17). For section immunostaining, postnatal mice were anaesthetized and perfused with phosphate-buffered saline (PBS, pH 7.4) through the right ventricle of the heart. The lung was inflated with 0.5% paraformaldehyde (PFA; P6148, Sigma) in PBS at 25 cm of H_2_O pressure. The lungs were then dissected and fixed with 0.5% PFA in PBS for 3-6 hr. For embryonic experiments, timed embryos were harvested from timed pregnancies and the trunks were separated along the ventral midline before being fixed in PBS with 0.5% PFA for at least 3 hr on a rocker at room temperature. For section staining, individual postnatal lobes were cryoprotected with 20% sucrose with 1/10 volume optimal cutting temperature medium (OCT; 4583, Tissue-Tek) in PBS and whole embryonic lungs were cryoprotected at 20% sucrose in PBS overnight on a rocker at 4°C. After cryoprotection, the samples were embedded in OCT. 10 µm OCT sections were washed with PBS, and blocked with PBS plus 0.3% Triton X-100 and 5% normal donkey serum (017-000-121, Jackson ImmunoResearch). Sections were then incubated with primary antibodies diluted in PBS plus 0.3% Triton X-100 in a humidity chamber at 4°C overnight. The sections were washed with PBS in a coplin jar for 30 min and incubated with secondary antibodies (1:1000, Jackson ImmunoResearch or Invitrogen) and 4′,6-diamidino-2-phenylindole (DAPI, 0.5 μg/ml final concentration) diluted in PBS plus 0.3% Triton X-100 at room temperature for 2 hr. The sections were washed again with PBS for 1 hr, then mounted with Aquapolymount (18606, Polysciences).

For whole-mount immunostaining, lobe strips or whole embryonic lung lobes were blocked with PBS plus 0.3% Triton X-100 and 5% normal donkey serum for 1 hr. After blocking, samples were incubated with primary antibodies diluted in PBS plus 0.3% Triton X-100 on a rocker at 4°C overnight. The next day, samples were washed for 3 hr with PBS plus 1% Triton X-100 and 1% Tween-20, and then incubated with secondary antibodies and DAPI in diluted in PBS plus 0.3% Triton X-100 on a rocker at 4°C overnight. The final day, the samples were washed for 3 hr with PBS plus 1% Triton X-100 and 1% Tween-20, then fixed with 2% PFA for 2 hr. To minimize experimental variations, littermate control and mutant lungs were embedded in the same OCT block or stained in the same tube. Images were captured on an optical projection tomography (OPT) microscope (Bioptonics) or confocal microscopes (FV1000, Olympus; A1plus, Nikon). Movies of 3D lungs were generated using Imaris (Bitplane).

### Whole mount in situ hybridization

Whole mount in situ hybridization was performed as previously described with minor changes (11). In brief, embryos from timed pregnancies were dissected in PBS and fixed in 0.5% PFA in PBS for 4-6 hr at room temperature on a rocker while genotyping was performed. Lungs were washed with diethylpyrocarbonate-treated PBS (DEPC-PBS) and dehydrated with methanol and stored at −20 °C. Samples were rehydrated in a methanol/DEPC-PBS gradient, washed with DEPC-PBS, and permeabilized in DEPC-PBS with 10 µg/ml protease K for 10 min for E11 lungs or 15 min for E15 lung lobes. Samples were re-fixed with 4% PFA and 0.25% glutaraldehyde for 20 min. The samples were then washed with DEPC-PBS and a rinsing solution (500 µl 4×SSC, 50% formamide, 0.1% Tween-20). Samples were blocked for 1 hr at 62°C in a slide oven in a prehybridization solution (50% formamide, 4×SSC, 1×Denhardt’s solution, 250 µg/µl salmon sperm DNA, 250 µg/µl yeast tRNA, 50 µg/ml heparin, 0.1% Tween-20). The samples were subsequently incubated at 62°C in a slide oven overnight with 1 µg/ml riboprobes diluted in 1 ml of hybridization solution (prehybridization solution with 10% >500 kDa dextran sulfate). The next day, the samples were washed with 50% formamide with 4×SSC and 0.1% Tween-20 at 65°C twice for 1 hr, incubated in 100 µg/ml RNase A diluted in 100 mM Tris (pH 7.5) with 500 mM sodium chloride and 0.1% Tween-20 at 37°C for 1 hr, washed in 50% formamide with 2×SSC and 0.1% Tween-20 at 65°C three times for 2 hr and then washed again overnight in a water bath. The following day, samples were washed in TBST (100 mM pH 7.5 Tris, 150 mM sodium chloride, 0.1% Tween-20) twice for 10 min, followed by blocking with TBST+5% normal sheep serum for 1 hr. An anti-digoxigenin alkaline phosphatase Fab fragment (1:2000, Roche, 11093274910) was then added to TBST for 2 hr followed by TBST washes four times for 2 hr and an overnight wash in TBST with 0.5 mg/ml levamisole (L9756, Sigma) at room temperature. On the last day, samples were washed with TBST for 30 min twice. The samples were then followed by detection of alkaline phosphatase activity with Nitroblue Tetrazolium and 5-bromo-4-chloro-3′-indolylphosphate (Roche, 11681451001) in the alkaline phosphatase reaction buffer (100 mM pH 9.5 Tris, 100 mM sodium chloride, 5 mM magnesium chloride). Following the alkaline phosphatase reaction, samples were fixed in 4% PFA in PBS for 2-4 hr, cleared via a 25%-50%-75% glycerol gradient, and mounted on a depression well slide for imaging.

Digoxigenin-labeled riboprobes (Roche, 11277073910) were transcribed with a T7 RNA polymerase (18033-019, Invitrogen) using cloned cDNAs or PCR products as templates. To minimize variation across experiments, littermate controls and mutants were processed in the same tube for each riboprobe throughout the whole experiment. Images were taken on an upright microscope (BX60, Olympus).

### OPT microscopy

OPT mounting and microscopy was performed as previously described (11, 17) with minor modifications. After whole embryonic lung immunostaining, samples were cleaned of debris on a stereomicroscope. Lungs were then washed and embedded in 1.5% low melting agarose (16520-100, Invitrogen) dissolved in water. After the agarose set, samples were cut and super glued onto the imaging stage. After 30 min of curing, mounted samples were dehydrated in two passages of methanol over two days, and 2 passages of BABB (one part benzyl alcohol (24122, Sigma) and two parts benzyl benzoate (B6630, Sigma)) over an additional day. Samples were imaged using the Bioptonics 3001 OPT scanner at standard resolution (512 pixels), according to the manufacturer’s instructions. Image stacks were reconstructed using the NRecon Reconstruction software (BrukermicroCT, BE) and visualized using maximal intensity projection (MIP) with the Bioptonics Viewer software (Bioptonics). Using ECAD and SOX2, lateral branch tips between branch lineage L1.A1 and L6 were counted as published (17).

### Cell dissociation and FACS for scRNA-seq

Cell dissociation and FACS was performed as previously described (22). Embryonic mouse lobes were dissected free of extra-pulmonary airways in iced Leibovitz media (Gibco, 21083), minced with forceps, and digested in Leibovitz with 2 mg/mL Collagenase Type I (Worthington, CLS-1, LS004197), 2 mg/mL Elastase (Worthington, ESL, LS002294), 0.5 mg/mL DNase I (Worthington, D, LS002007) for 20 min at 37 °C. The tissue was mechanically titrated halfway through digestion. Fetal bovine serum (FBS, Invitrogen, 10082-139) was added to a final concentration of 20% and the solution was triturated until homogenous. The sample was transferred to the cold room on ice and was filtered with a 70 µm cell strainer (Falcon, 352350), and spun down at 2,040 rcf for 1 min. The cells were resuspended in red blood cell lysis buffer (15 mM NH_4_Cl, 12 mM NaHCO_3_, 0.1 mM EDTA, pH 8.0) for 3 min twice, washed with Leibovitz with 10% FBS, and filtered into a 5 ml glass tube with a cell strainer cap (Falcon, 352235). The cells were then incubated with PE/Cy7-conjugated rat anti-CD45 (CD45, 1:250, 103114, BioLegend), A647-conjugated rat anti-intercellular adhesion molecule 2 (ICAM2, 1:250, A15452, Invitrogen), Alexa488-conjugated rat anti-E-cadherin (ECAD, 1:250, 53-3249-80, eBioscience) at a concentration of 1:250 for 30 min, spun down at 2,040 rcf for 1 min, washed for 5 min and resuspended with Leibovitz with 10% FBS. The sample was refiltered and incubated with SYTOX Blue (Invitrogen, S34857), then sorted on a BD FACSAria Fusion Cell Sorter. After exclusion of dead cells, CD45 and ICAM2 negative cells were selected and from those ECAD positive cells were collected and processed through the Chromium Single Cell Gene Expression Solution Platform (10X Genomics) using the Chromium Single Cell 3’ Library and Gel Bead Kit in accordance with the manufacturer’s user guide (v2, rev D). The libraries were sequenced on an Illumina NextSeq500 using a 26×124 sequencing run format with 8 bp index (Read1). Raw data have been deposited in GEO under the accession number GSE144170. AT1 and AT2 gene lists are from the following dataset: GSE124325 (26).

### Single-cell RNA-seq data analysis

scRNA-seq output was processed using Cell Rangers “cellranger count” and “cell ranger aggr”. Downstream analyses were carried out using the Seurat R package (38–40) and Monocle R Package (41–43). We used Seurat 3.1.2 to normalize and scale the aggregated dataset and separate out the epithelial cells from the contaminating mesenchymal cells by marker expression of ECAD. In the epithelial subset, we further defined cell populations by using markers for progenitors (*Sox9*), their progeny (*Sox2*), and neuroendocrine cells (*Sox2* and *Ascl1*). We then identified differentially expressed genes for each cell type between control and mutant and generated volcano plots to visualize the up and down regulated genes.

For trajectory analysis, Monocle 2.8.0 (R package) was used on the control and mutant SOX9 cells. The dataset was formed with a negative binomial distribution, loading the phenoData and featureData from Seurat. The data was reduced by DDRTree, and the trajectory was generated using the top 500 variable genes. Differentially expressed genes from Seurat were graphed on the trajectories to visualize the trend in expression changes. Signature genes for AT1, AT2, and progenitor cells were used for metagene analysis using Seurat’s module score function. scRNA-seq data from the Lef/Tcf mutant were plotted against the published bulk RNA-seq data from the *Ctnnb1* mutant (11).

## ACKNOWLEDGEMENT

We thank Dr. Hai-Hui Xue (University of Iowa) for sharing the *Lef1*^*LoxP*^ and *Tcf7*^*LoxP*^ mice, Dr. Elaine Fuchs (Rockefeller University) and Dr. Hoang Nguyen (Baylor College of Medicine) for sharing the *Tcf7l1*^*LoxP*^ mice, and Drs. Mario Capecchi and Melinda Angus-Hill (University of Utah) for sharing the *Tcf7l2*^*LoxP*^ mice. We thank Dr. Hoang Nguyen (Baylor College of Medicine) for sharing the TCF7L1 antibody. The University of Texas MD Anderson Cancer Center DNA Analysis Facility and Flow Cytometry and Cellular Imaging Core Facility are supported by the Cancer Center Support Grant (CA #16672). This work was supported by the University of Texas MD Anderson Cancer Center Start-up and Retention Fund, an Institutional Research Grant, and National Institutes of Health R01HL130129 (JC).

## AUTHOR CONTRIBUTIONS

KNGM and JC designed and performed research, and wrote the paper; HA provided the *Sox9*^*CreER*^ mice; all authors read and approved the paper.

---

**Supplemental Figure 1:**
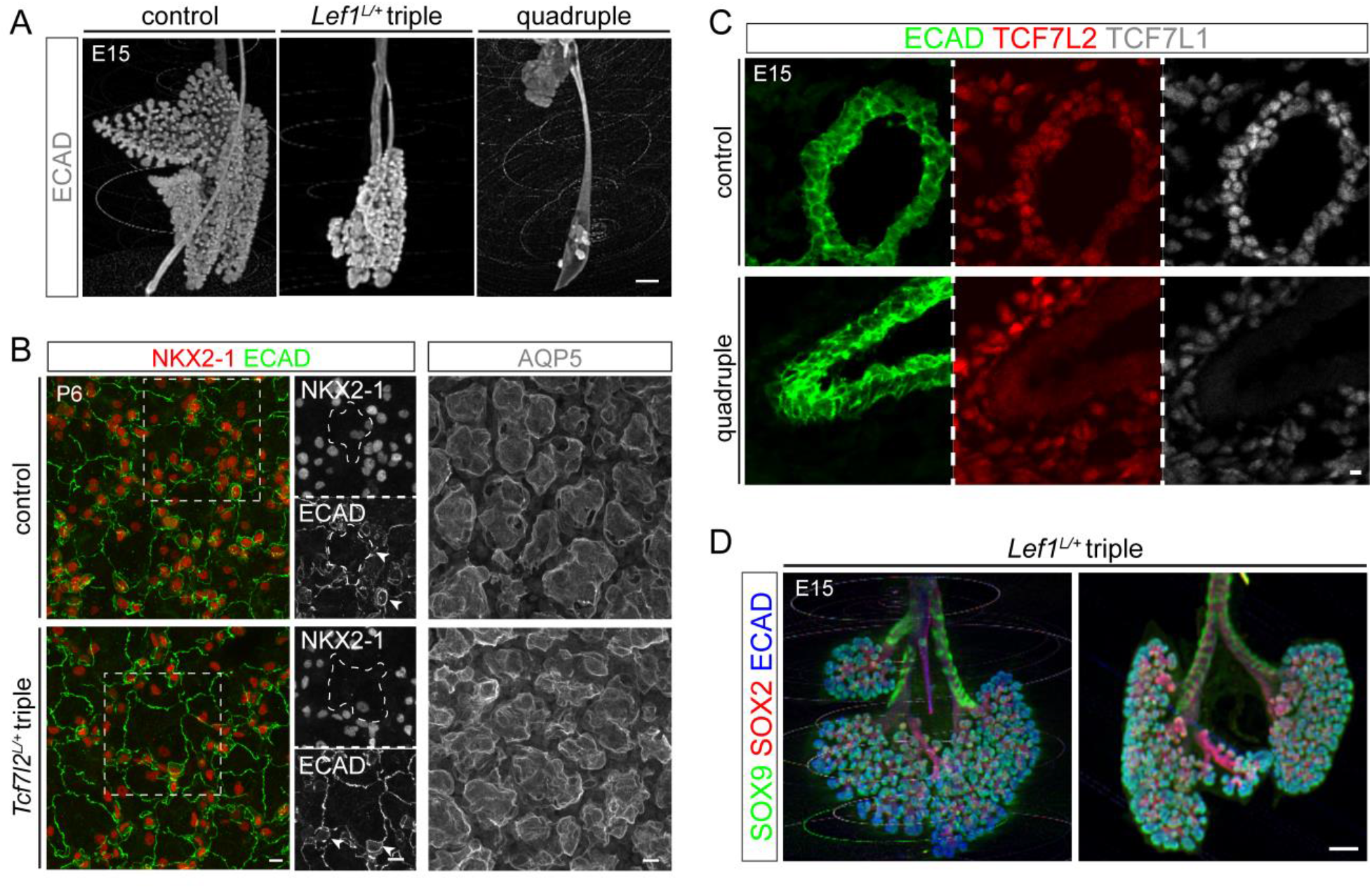
Additional phenotype characterization of pan-epithelial (*Shh*^*Cre*^) Lef/Tcf mutants. **(A)** Side view of OPT images shown tracheoesophageal fistula in *Lef1*^*L/+*^ triple mutant and quadruple mutant lungs. **(B)** En face confocal projection images of immunostained postnatal *Tcf7l2*^*L/+*^ triple mutant and littermate control lungs showing normal AT1 (dashes) and AT2 (arrowheads) differentiation and alveolar morphology (AQP5). **(C)** Confocal images of immunostained quadruple mutant and littermate control lungs showing loss of TCF7L1 and TCF7L2. LEF1 and TCF7 are normally expressed at a level too low to be informative. **(D)** OPT images of 2 *Lef1*^*L/+*^ triple mutant lungs with a normal number of lateral branch tips in Fig. 2D but defective branching. All images are representative of at least 2 biological replicates. Scale bars: 10 µm for confocal images and 250 µm for OPT images.

**Supplemental Videos: OPT images of lungs referred to in figure legends.**

**Supplemental Table 1: Differentially expressed genes for volcano plots in Fig. 4C.**

**Supplemental Table 2: Gene lists for Monocle branch heatmaps in Fig. 4F.**

**Supplemental Table 3: Signature genes for AT1, AT2, and progenitor cells for Fig. 4G.**

**Supplemental Table 4: Differentially expressed genes comparing AT1 versus AT2 cells using an E19 scRNA-seq dataset for Fig. 4G.**

**Supplemental Table 5: Comparison of quadruple mutant scRNA-seq and *Ctnnb1* mutant bulk RNA-seq.**

